# Lipopolysaccharide stimulation of RAW264.7 cells is a model for identifying novel clients of Hsc70

**DOI:** 10.1101/2020.05.15.098525

**Authors:** John F. Rakus, Nicholas R. Kegley, Alex J. Ashley, Michael A. Parsons, Megumi Takeuchi

## Abstract

Heat shock cognate protein 71 kDa (Hsc70, Hspa8, Hspa10, Hsp73) is a member of the heat shock protein 70 kDa family of molecular chaperones. These chaperones function to aid the correct folding of client proteins using an ATP-dependent mechanism. Though Hsc70 is accepted to be a constitutively expressed protein, it has a well-documented function in modulating the induction of the pro-inflammatory cytokine TNFα by the LPS/TLR4 pathway. In this work we attempt to identify protein clients of Hsc70 to gain insight into those which may be responsible for this regulatory effect. RAW264.7 cells were cultured in the absence or presence of 1 μg/mL LPS for 0 to 24 hours. Herein we describe of a large number of newly-categorized Hsc70 clients using immunoprecipitation and nanoLC-MS/MS and we validate several novel Hsc70/client interactions using co-immunoprecipitation. After performing immunoprecipitation using a commercially available antibody, eluted fractions were proteolytically digested either immediately after immunoprecipitation or after separation by SDS-PAGE and sequence analyzed using nanoLC-MS/MS with a Q-Exactive Plus triple-quadrupole/orbitrap mass spectrometer. Using these methods, 292 total unique protein hits were identified with high confidence; 34 of which were only detected in LPS-treated cells.

## Introduction

The function of chaperone proteins is to assist the proper folding of client proteins in order to ensure that the client is fully functional.^1^ This is usually an ATP-driven process due to the large thermodynamic commitment necessary to synthesize a full-length polypeptide and the deleterious effects that can occur by improperly folded proteins in a cell. The heat shock protein 70 kDa (Hsp70) family contain bi-domain chaperone proteins with a general architecture consisting of an N-terminal ATPase domain and a C-terminal substrate-binding domain to catalyze conformational changes in the client upon hydrolysis of ATP.^2^ The chaperones of this family tend to display specificity towards hydrophobic regions of protein structure which are normally buried in a properly folded polypeptide.^3^ As the name suggestions, heat shock proteins are often induced upon a cellular stressor, but at least one member – heat shock cognate protein 71 kDa (Hsc70; also Hspa8, Hspa10, Hsp73) – is thought to be a constitutively expressed chaperone.^4^

Few validated clients of Hsc70 are known but some include thrombospondin-1,^5^ XIAP,^6^ HMGB1,^7^ stathmin,^8^ heat-shock factor 1^9^ and heme-regulated inhibitor.^10^ In addition to its chaperone function, Hsc70 is involved in clathrin-mediated vesicle coating,^11,12^ protein translocation^13,14^ and lysosomal targeting^15^ as well as many other cellular processes and is implicated in many diseases (reviewed in Ref. 1). Substantial work has demonstrated that Hsc70 has a role in modulating the lipopolysaccharide (LPS)/toll-like receptor 4 (TLR4) pathway in cardiac myocytes.^16–18^ This implies that Hsc70, or by extension one of its clients, is involved in the regulation of this pro-inflammatory pathway which stimulates the release of tumor necrosis factor α (TNFα), the central cytokine of inflammation. Further work in the mouse RAW264.7 macrophage-like cell line has indicated that this TNFα-inducing effect is enhanced in the presence of a C-mannosylated thrombospondin-1-derived peptide.^19^ Later work indicated that a C-mannosyltryptophan moiety – a modification commonly observed in thrombospondin type-1 repeat (TSR) domains – was a requirement for Hsc70-facilitated TNFα secretion and activity.^5^ The demonstrated effect of Hsc70 on the LPS/TLR4 pathway, particularly through an interaction with C-mannosyltryptophan, suggests that cataloging the possible clients of this chaperone can serve to identify system-specific, biologically-relevant interactions of Hsc70 and may determine whether thrombospondin-1 is a valid regulator of TNFα in the LPS/TLR4 pathway, or if another C-mannosylated client is responsible.

Currently there is limited knowledge about the range of targets of Hsc70 activity. Intrigued by the observations of Muroi *et al.* and Ihara *et al.*, we decided to replicate their original experiments in RAW264.7 cells treated with 1 μg/mL LPS with the intention of using Hsc70 as a probe to target potential clients, particularly those whose expression may be a consequence of LPS treatment. In order to determine the identity of Hsc70 clients, we devised an immunoprecipitation strategy coupled to mass spectrometry to determine proteins bound by Hsc70 in RAW264.7 cells. We used a commercially available monoclonal antibody to target the endogenously expressed murine Hsc70 (mHsc70). Two sample preparation strategies were employed. In the first the antibody was non-covalently bound to Protein G-conjugated magnetic beads and eluted with SDS-sample buffer. Elutions were separated by SDS-PAGE and all visible silver-stained bands were excised and proteolytically digested with trypsin and chymotrypsin. We then performed nanoLC-MS/MS to sequence peptides derived from those bands. In the second complementary method, the anti-mHsc70 antibody was crosslinked to Protein-G-conjugated magnetic beads using disuccinimidyl suberate and eluted by heating at low pH. The elutions were pH neutralized, proteolytically digested in solution with trypsin and then subjected to nanoLC-MS/MS. After conducting an isotype-control antibody immunoprecipitation to discard candidates that may be interacting with the antibody/bead/Protein G complex, we proceeded to classify the remaining sequences as potential Hsc70 clients. Combined, we were able to detect 292 separate, unique protein sequences based on the presence of at least two peptides with greater than 95% confidence. This included 34 proteins only identified in LPS-treated samples, 141 only in non-LPS-treated samples and 117 which were in both. Using the MS/MS results as a guide, we further identified receptor expressed in lymphoid tissues (RELT) as previously unknown clients of Hsc70. These results imply that many of the proteins identified by this method engage in biologically relevant associations with mHsc70.

## Results and Discussion

We initially validated our method by performing immunoprecipitation of mouse Hsc70 (mHsc70) from RAW264.7 cells treated with or without 1 μg/mL LPS. This LPS concentration is well above what will initiate the TLR4 pathway,^20^ however we reasoned that adhering to the conditions originally set forth by Muroi *et al.*, and Ihara *et al.* necessitated using this amount^5,19^ Growth media began to turn yellow after approximately six hours and a large fraction of cells detached from the flask surface. Robust secretion of TNFα to the media and strong but comparatively lesser secretion of IL-6 were observed only in LPS-stimulated cells confirming activation of the LPS/TLR4 pathway (Figure S-1).

Initially, we chose to utilize a traditional in gel strategy to identify proteins which might interact with mHsc70 under these growth conditions. The SDS-buffer eluents for each condition were separated by SDS-PAGE and silver stained (Figure 1A). We initially intended to investigate mHsc70 interactions up to 48 hours post-LPS, however due to substantial protein degradation those samples were not pursued further. All visibly distinguishable bands were excised and reduced with TCEP followed by alkylation of cysteines with iodoacetamide. The polypeptides were then proteolytically digested with trypsin and chymotrypsin. Presence of mHsc70 for each condition was verified by mass spectrometric detection in the bands at approximately 70 kDa as indicated in Fig. 1A and mHsc70 was confirmed with greater than 50% sequence coverage in these bands (Fig. 1B). Peptides from mHsc70 were validated with high confidence, as demonstrated by three fragment ion series with a near complete set of either b- or y-ions and a substantial number of matched ions (Fig. 1C-1E). Hsc70 did not elute when this immunoprecipitation method was performed with either non-conjugated beads or isotype control antibody-conjugated magnetic beads (Figure S-2). We acknowledge that the harsh lysis conditions we used would likely result in release of organelle-resident proteins such as those in the mitochondrion. Heat-based cell lysis can be adopted for proteomic research^21–23^ and Hsc70 has been shown to be involved in delivery of cytosolic proteins to the nucleus, mitochondrion and endoplasmic reticulum.^24,25^ Due to this interaction and the observation that Hsc70 recognizes mitochondrial aspartate aminotransferase but not cytosolic,^26,27^ we reason that our protocol is consistent with the expectation of producing biologically relevant interactions.^28–31^ Based on these data we proceeded to analyze all 115 visible bands between 10 and 250 kDa in Figure 1A.

**Figure 1.**
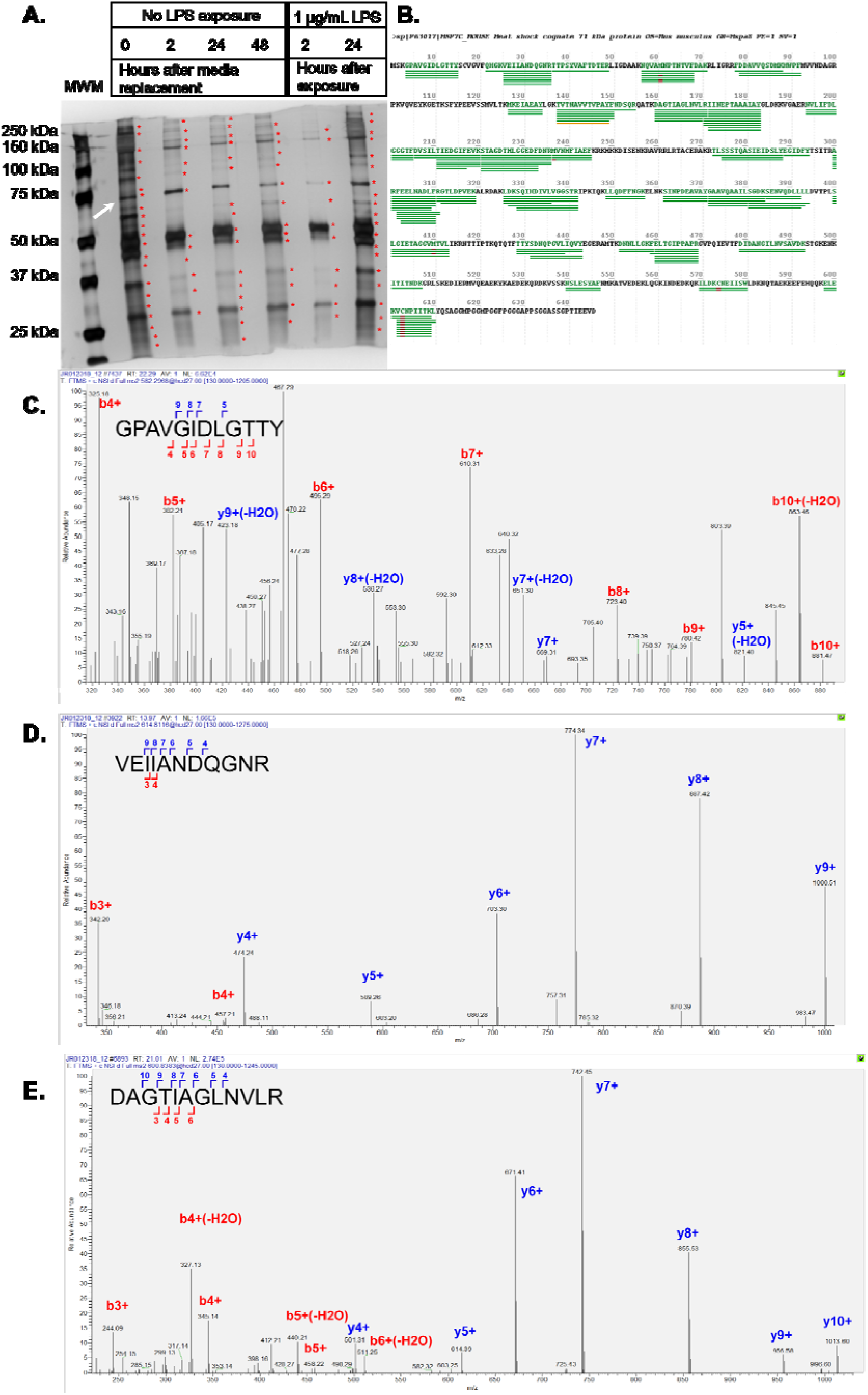
Immunoprecipitation of mHsc70 from the RAW264.7 cell line. A. Silver-stained SDS-PAGE of eluted anti-Hsc70 immunoprecipitations from RAW264.7 cells grown in serum-depleted conditions with or without 1 μg/mL LPS. Bands indicated with 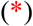 were excised and analyzed for mass spectrometry. B. Sequence coverage map of mHsc70 isolated from the 0 hour control lane (specific band indicated with a white arrow in panel A). Red underlines indicate carbamidomethyl-cysteine residues; green lines indicate peptides which were identified by MS/MS. C. Representative fragment ion series from the peptide G^4^PAVGIDLGTTY^15^ (observed m/z 582.298, observed mass (+H) 1163.558 Da, +2 charge, −5.4 ppm mass error, Byonic score 224.3), D. V^26^EIIANDQGNR^37^ (observed m/z 614.812, observed mass (+H) 1228.616 Da, +2 charge, −10.0 ppm mass error, Byonic score 361.3) and E. D^160^AGTIAGLNVLR^171^ (observed m/z 522.286, observed mass (+H) 1043.595 Da, +2 charge, −7.8 ppm mass error, Byonic score 202.5).

In total, 1028 unique protein sequences were initially identified from the in gel digestions based on the minimum of one peptide identified in Byonic searches (Table S-1). Byonic was used specifically to identify glycosylated peptides, particularly those which might contain C-mannosyltryptophan. To refine the data set, “low quality” peptide hits were removed using criteria described in Wong *et al.*^7^ by setting a threshold of two unique peptides each at >95% confidence to qualify as a protein “hit,” leaving 520 sequences remaining.

We decided to refine our immunoprecipitation strategy by performing a complementary in solution proteomic analysis with biological replicates of those analyzed by in gel digestion. The immunoprecipitation protocol was also modified by covalently crosslinking the anti-mHsc70 primary antibody to the Protein G-Dynabead complex. In these samples, mHsc70 binding partners were eluted by heating in a low-pH buffer as opposed to reducing SDS-PAGE loading buffer in order to maintain optimal conditions for direct in solution proteolysis after neutralization. The proteins were reduced, alkylated and trypsin digested then subjected to the same nanoLC-MS/MS protocol, Byonic searches and thresholding as described for the in gel-digested samples leaving 699 total protein targets identified (Table S-1). Combining the result of both immunoprecipitations, 1023 total unique protein hits met the threshold to be considered a valid hit. In order to ensure that these mHsc70 client candidates were the result of specific interactions between the client and chaperone, the samples described above were also analyzed by the in solution method described above using a covalently crosslinked isotype control antibody conjugated to the Protein G-Dynabead complex (Table S-2).^32^ After thresholding, 292 sequences unique to mHsc70 immunoprecipitation remained. Of these, 117 sequences were identified in both LPS-treated and control samples, 141 were identified only in control samples and 34 were only identified in LPS-treated cells (Table S-3). Data are available *via* ProteomeXchange with identifier PXD016206.^33,34^

To assess the biological relevance of these clients, the cohort of 292 proteins from both digestions were analyzed for functional similarity enrichment using DAVID (Database for Annotation, Visualization and Integrated Discovery) ver. 6.8 (david.ncifcrf.gov).^35,36^ Only clusters with DAVID enrichment scores greater than 1.00 and containing a GeneOntology entry with p < 0.005 were considered (Table S-4). We deduce that any functional clusters identified with these criteria are likely to be preferentially enriched through Hsc70 interactions above a normal background proteome. We observed substantial enrichment of known Hsc70 cochaperones including several Hsp40 proteins and TCP-1 confirming Hsc70-specific interactions in these immunoprecipitations. The previously identified mHsc70 client HMGB1 was also detected in six of the eight separate samples analyzed by the in solution method. Additionally, enrichment of proteins related to RNA binding, unfolding protein binding, carbon metabolism and mitochondrial function were observed.

We then proceeded to analyze the LPS-only and control-only cohorts from both immunoprecipitation experiments with DAVID. The LPS-treated cluster was enriched in RNA-binding and mitochondrial inner membrane proteins; the latter consistent with the observations that LPS stimulation of macrophages initiates mitochondrial destabilization (Table 1, Table S-4).^37^ The control clusters were substantially enriched in metabolic proteins related to carbon metabolism, cytoskeletal binding and adherence, nucleotide binding, the mitochondrion and unfolded protein binding.

**Table 1.**
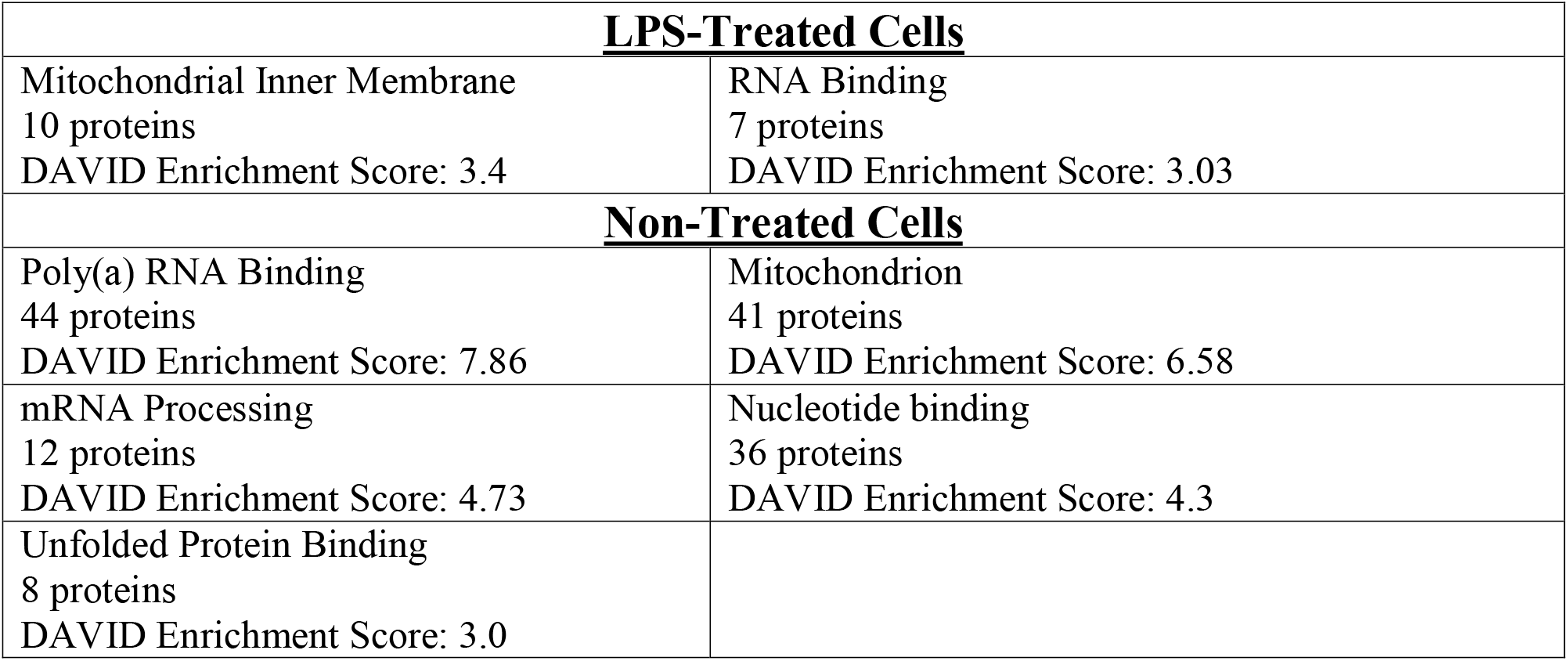
Listing of functional identifications for high-confidence protein hits as organized by function by DAVID. The DAVID enrichment score for each cluster is indicated as well as the gene ontology (GO) or Kyoto Encyclopedia of Genes and Genomes (KEGG) pathway entry for each cluster. Complete list of entries in each cluster in Table S-4.

| **<u>LPS-Treated Cells</u>** |  |
| --- | --- |
| Mitochondrial Inner Membrane<br>10 proteins<br>DAVID Enrichment Score: 3.4 | RNA Binding<br>7 proteins<br>DAVID Enrichment Score: 3.03 |
| **<u>Non-Treated Cells</u>** |  |
| Poly(a) RNA Binding<br>44 proteins<br>DAVID Enrichment Score: 7.86 | Mitochondrion<br>41 proteins<br>DAVID Enrichment Score: 6.58 |
| mRNA Processing<br>12 proteins<br>DAVID Enrichment Score: 4.73 | Nucleotide binding<br>36 proteins<br>DAVID Enrichment Score: 4.3 |
| Unfolded Protein Binding<br>8 proteins<br>DAVID Enrichment Score: 3.0 |  |

We were also interested in identifying novel clients of mHsc70 with potential relation to the LPS/TLR4 pathway, which led us to focus on Receptor Expressed in Lymphoid Tissues (RELT; TNFRSF19L), which was originally identified by one peptide with medium-high confidence in the elution from cells after 24 hours post LPS treatment. RELT is a member of the TNFα receptor superfamily but its activating ligand is unknown.^38,39^ In order to probe a possible relevance between mHsc70 and RELT, endogenous RELT was successfully immunoprecipitated from RAW264.7 cells in eluents which included mHsc70, identified by both immunoblot and mass spectrometry. (Figure 2). The mHsc70 peptides D^160^AGTIAGLNVLR^171^ (Fig. 2B) – which was also detected in the elution for Hsc70 immunoprecipitation (Fig. 1E) – and F^302^EELNADLFR^311^ (Fig. 2C) demonstrate the presence of mHsc70 in the RELT immunoprecipitation elution.

**Figure 2.**
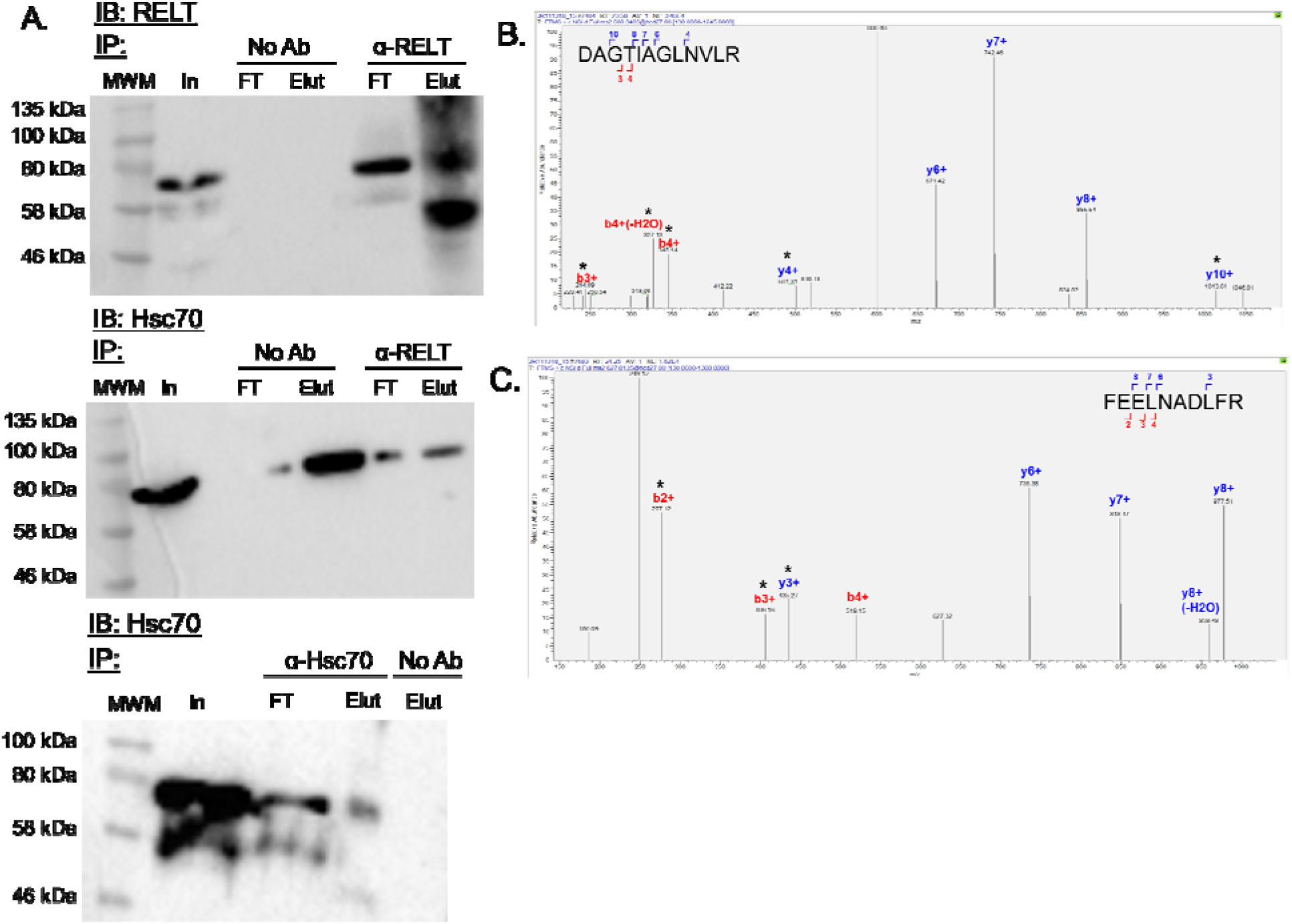
Interaction of mRELT with mHsc70. A. Immunoblot of RELT immunoprecipitations (IP) from RAW264.7 cells grown under standard culture conditions. Hsc70 and RELT were immunoblotted (IB) separately and proteins are indicated with arrows. MWM: molecular weight marker; No Ab: elution of non-antibody-conjugated Protein G beads; FT: flow-through of lysate after antibody/bead complex addition; elution: 70 °C, 10 minutes in SDS-PAGE buffer. Images were captured with a Bio-Rad Molecular Imager ChemiDoc XRS+ using Image Lab 5.2.1. For each blot, the image was captured twice, first using white epi-illumination to capture the Broad Range Color Protein Standard (New England Biolabs) followed by chemiluminescence to capture the sample bands. The saved imaged where overlaid in Image Lab 5.2.1 using the Merge Image function. B. Representative fragment ion series of the D^160^AGTIAGLNVLR^171^ (observed m/z 600.340, observed mass (+H) 1199.673 Da, +2 charge, −0.8 ppm mass error, Byonic score 376.5)and C. F^302^EELNADLFR^311^ (observed m/z 627.314, observed mass (+H) 1253.620 Da, +2 charge, +3.0 ppm mass error, Byonic score 319.4) peptides. Isotopic distributions for peaks labeled with (*) were unable to be extracted from the data results.

## Conclusions

This project was aimed towards identifying client proteins of mHsc70 with a particular interest in distinguishing how activation of the LPS/TLR4 pathway in RAW264.7 macrophages affects interactions with those clients. These cells were challenged with either 1 μg/mL of LPS or non-treatment control over time periods of up to 48 hours and cell lysates were isolated in order to analyze cytosolic and membrane-derived proteins. Immunoprecipitation of mHsc70 was successfully achieved and in gel and in solution proteolytic digests and were utilized as the means to identify Hsc70 co-eluted proteins. This revealed 292 co-precipitated proteins identified by two unique peptides at a confidence interval >95% each, 117 of which were detected by both treatments. The fact that eighty percent of the identified protein targets were only detected with one separation technique is indicative of the necessity to use multiple complementary strategies to obtain biologically relevant proteomic information. Of the proteins identified, 34 were only observed in LPS-treated samples and 141 were only observed in the non-challenged cells (Table S-1). Our informatics analysis revealed that co-precipitated proteins involved in RNA binding, carbon metabolism and protein folding. These co-precipitated proteins include known Hsc70 cochaperones from the Hsp40 as well as the previously reported Hsc70 client HMGB1. The presence of these known Hsc70 binding partners in our result is indicative that we are reporting authentic Hsc70 client candidates.

In the design of this experiment, we had hoped to be able to identify potentially C-mannosylated peptides in order to gain insight into identifying the protein(s) responsible for the effect observed when C-Man-WSPWS is provided to LPS-treated RAW264.7 cells. Our mass spectrometry results, however provided few high-confidence peptides to indicate C-mannosyltryptophan in the WXX(W/C) consensus sequence. This should not necessarily be taken as an indication that this modification is absent in the cohort identified in this work. More than half of the unique protein sequences which provided at least one peptide hit only provided one hit and many were of very low confidence with poor signal intensity and MS/MS fragmentation. C-mannosylated proteins may be co-eluting with mHsc70, but C-mannosyltryptophan-containing peptides were not detected in our MS searches. Further, we accept that the our experimental design does not necessary provide an exhaustive catalogue of every possible mHsc70 client and due to our focus in capturing peptides from a broad range of proteins, authentically C-mannosylated peptides may have escaped our detection.

Lastly, we were interested in validating novel mHsc70 complexes which may be involved in the LPS response. To that end, we observed that RELT, an orphan TNFα receptor family member, was co-precipitated in LPS-treated cells. Immunoprecipitation of endogenous RELT from RAW264.7 cells exposed to LPS for twenty-four hours indicates both RELT and mHsc70 are present. This is indicative of the possibility that RELT is a client of mHsc70 and that mHsc70 may function as a ligand and may affect RELT activity. In summary, we have identified a large swath of previously unknown Hsc70 client proteins including distinct cohorts of client proteins observed based on the presence or absence of LPS. This has led to the confirmation of previously reported mHsc70-interacting proteins and known mHsc70 clients indicating that this chaperone may influence many other biological processes. Our identification of RELT as a potential mHsc70 client despite being originally detected with a mass spectrum below our threshold implies that we may be able to use less stringent criteria in order to validate mHsc70 interactions by immunoprecipitation.

## Experimental

### Materials

All chemicals used for this experiment were of the highest available molecular biology grade. Lipopolysaccharides (LPS) from *Escherichia coli* O55:B5 was purchased from Millipore Sigma (Burlington, MA). Primary antibodies against mHsc70 and RELT; and HRP-conjugated goat anti-mouse-IgG, goat-anti-rabbit-IgG and goat-anti-rat-IgG secondary antibodies were purchased from ThermoFisher Scientific (Waltham, MA). The rat-IgG1κ isotype control antibody was purchased from Abcam Inc. (Cambridge, United Kingdom). 680RD-conjugated goat anti-mouse-IgG and goat anti-rat-IgG and 800CW-conjugated goat anti-human-IgG secondary antibodies were purchased from LI-COR Biotechnologies (Lincoln, NE).

### Cell culture

RAW264.7 were a generous gift from Dr. Fikri Avci (University of Georgia) and originally purchased from the American Type Culture Collection (Manassas, VA). Cells were cultured generally at 37 °C in 5% CO_2_ in high glucose Dulbecco’s Modified Eagle’s Medium (DMEM) supplemented with 5% fetal bovine serum (FBS) and penicillin/streptomycin (pen/strep). Cells were passaged at a 1:6 ratio and no cells past the tenth passage were used for analysis.

### LPS experiment

Cells were first acclimated to serum-free media by incubation in DMEM and pen/strep for 24 hours prior to introduction of LPS. To acclimate, media was aspirated and the cells gently washed once with Dulbecco’s phosphate buffered saline (PBS) before replacement of an equal volume of serum-free media. After the acclimation period, the media was aspirated and replaced with an equal volume of serum-free media (control conditions) or serum-free media supplemented with 1 μg/mL LPS (treatment conditions) to replicate conditions from Muroi *et al*.^19^ and Ihara *et al.*^5^ Cells were incubated and collected at 1, 2, 6, 24 and 48 hours post-stimulation. Four 10 cm cell culture dishes per condition were used to isolate protein.

### Sample preparation

Conditioned media was aspirated and flash frozen with liquid nitrogen. Cells were gently washed twice with Dulbecco’s PBS and washes were pooled with the conditioned media. One mL of Dulbecco’s PBS with protease inhibitors was added to the growth flask and cells were gently scraped and pipetted. The cells were centrifuged for five minutes at 1000 rpm followed by the addition of one mL Tris buffered saline (TBS) with 1% Triton X-100 and protease inhibitors. Cell lysates were placed at 37 °C for one hour with 15 seconds sonication every 15 minutes.^21,22^ Lysates were then centrifuged ten minutes at 12,700 rpm and the supernatants were collected.

### Immunoprecipitation experiments

2 μg of anti-mHsc70 primary antibody (1B5, ThermoFisher) was resuspended in 200 μL PBS + 0.02% Tween-20 (PBS-T). 50 μL of Protein G-conjugated Dynabeads (ThermoFisher) were washed once with PBS-T. The antibody was then transferred to the Dynabeads and incubated at room temperature for ten minutes in gentle agitation. The beads were rinsed gently with PBS-T three times. 500 μg of a protein lysate, as determined by BCA assay (ThermoFisher), was added to the bead tubes, mixed, and set to incubate at 4 °C overnight while rotated gently. The supernatant was then aspirated, beads gently rinsed three times with PBS-T and proteins dissociated from the antibody/bead complex by incubation at 70 °C for ten minutes in SDS-PAGE running buffer (200 mM Tris, pH 6.8, 40% glycerol, 80.0 mg/mL SDS, 4 mg/mL bromophenol blue, 8 μL/mL β-mercaptoethanol). Samples for in gel mass spectrometry analysis were separated on SDS-PAGE using either 10% pre-cast Tris/glycine or 5%/10% Tris/glycine gels by running samples at 60 V for 45 minutes and 180 V for approximately 60 minutes. The gels were subsequently stained using the Silver Stain for Mass Spec Kit (ThermoFisher).

In order to minimize interference from the anti-mHsc70 antibody in immunoprecipitations, the experiment was repeated using covalently-crosslinked Protein G Dynabeads. Prior to incubation of antibody/bead complex with lysates, the primary antibody was crosslinked to Protein G using disuccinimidyl suberate (DSS) by incubating the antibody/bead complex in 625 μM DSS at room temperature for 30 minutes followed by quenching with 1.0 M Tris (pH 7.5) for 15 minutes at room temperature then rinsing the beads once with 200 μL PBS-T. Lysates were then added to the antibody/bead complex, incubated, washed and eluted in 50 μL ThermoFisher IP Elution buffer at 70 °C for ten minutes. This protocol was also used for the isotype-control immunoprecipitations. Immunoprecipitation for RELT followed a similar protocol. However prior to incubation of antibody/bead complex with lysates, the primary antibody was crosslinked to Protein G using DSS as described above. Lysates were then added to the antibody/bead complex, incubated, washed and eluted in SDS-PAGE sample buffer.

### Mass spectrometry

Excised gel bands were reduced in 100 μL of 50 mM Tris (pH 6.8) with 20 mM tris(2-carboxyethyl)phosphine (TCEP) at 100 °C for five minutes. After bands cooled to room temperature, the buffer was aspirated and 150 μL of 33.3 mM iodoacetamide in 50 mM Tris (pH 6.8) was added and reacted at room temperature in the dark for 30 minutes. The buffer was aspirated and gel bands were washed with one mL of 50% methanol in 20 mM diammonium phosphate (wash buffer) for one hour at room temperature then overnight at 4 °C in the same. Afterward, a sample was washed three additional times for 30 minutes each at room temperature in wash buffer. One mL of mass spectrometry-grade acetonitrile was added and gel samples were rotated at room temperature for 30 minutes. Acetonitrile was aspirated and bands set to dry. Proteins were proteolytically digested in 30 μL of 20 mM diammonium phosphate with 2 mM calcium chloride, trypsin (16.7 ng/μL) and chymotrypsin (16.7 ng/μL) for 5.5 hours at 37 °C. Samples were then sonicated in a water bath for 20 minutes, the supernatant was saved and the gel pieces were sonicated for an additional 20 minutes in 5% formic acid. Supernatants were pooled and centrifuged through a 3 kDa MW cutoff filter then desalted with a C-18 ZipTip column. In solution digestions were reduced and alkylated as described above then incubated in 30 μL of 20 mM ammonium bicarbonate with 16.7 ng/μL trypsin at 37 °C overnight. Reactions were quenched by the addition of approximately 7 μL 5% formic acid and sonication for 20 minutes. Peptides were purified through the same protocol as described above. For all sample preparations, the resulting peptides were diluted to 25% acetonitrile and injected onto an Easy nLC 1000 HPLC system with reverse phase column ES801 (ThermoFisher) coupled to a Q-Exactive Orbitrap mass spectrometer (ThermoFisher). Separation of (glyco) peptides was performed with a binary gradient solvent system that consists of solvent A (0.1% formic acid in water) and solvent B (80% acetonitrile and 0.1% formic acid in water) with a constant flow rate of 300 nL/min. Spectra were recorded with a resolution of 70,000 in the positive polarity mode over the range of m/z 400-2,000 and automatic gain control target value was 1 × 10^6^. The ten most prominent precursor ions in each full scan were isolated for higher energy collisional dissociation-tandem mass spectrometry (HCD-MS/MS) fragmentation with normalized collision energy of 27%, and automatic gain control target of 1 × 10^5^, an isolation window of m/z 1.2 daltons, dynamic exclusion enabled, and fragment resolution of 17,500. Raw data files were analyzed using Proteome Discoverer ver. 2.1 (ThermoFisher) with Byonic ver. 2.10.5 (Protein Metrics, Cupertino, CA) as a module for automated identification of (glyco) peptides. All searches were performed on a library of the mouse proteome downloaded from the UniProt proteome database. Searches were conducted with a 10.0 ppm precursor tolerance, 20.0 ppm fragment tolerance and a maximum of four missed cleavages for trypsin/chymotrypsin in gel proteolytic digests with cleavages occurring C-terminal to lysine, arginine, tyrosine, phenylalanine, tryptophan or leucine residues. The following amino acid modifications were included in these searches: carbamidomethyl-cysteine (+57.021464 Da), oxidized methionine (+15.994915 Da), C-hexosyl-tryptophan (+162.052824 Da), deoxyhexosyl-serine/threonine (+146.057909 Da) and deoxyhexosyl-hexosyl-serine/threonine (+308.110732 Da). For in solution tryptic digests, the missed cleavages threshold was set to a maximum of four occurring C-terminal to lysine and arginine residues. All other search conditions remained the same.^40^

A parallel reaction monitoring (PRM) experiment was recorded with a resolution of 35,000 in positive polarity mode over the range m/z 300-2,000 and an automatic gain control target value was 3 × 10^6^. Targeted peptides were isolated for HCD-MS/MS fragmentation with normalized energy of 27%, and automatic gain control target of 2 × 10^5^, an isolation window of m/z 2.0 daltons, and fragmentation resolution of 17,500.

### Western blot

Samples for western blot were separated by SDS-PAGE using either 10% pre-cast Tris/glycine gel 5% stacking/10% separating Tris/glycine gels by running samples at 60 V for 45 minutes and 180 V for approximately 60 minutes. Separated proteins were transferred to 0.45 μm nitrocellulose paper at 0.50 A current for one hour in Towbin buffer [25 mM Tris, 192 mM glycine, 20% methanol (v/v)]. The blots were incubated in blocking buffer (5% non-fat dried milk in TBS with 0.02% Tween-20) at room temperature for thirty minutes. The primary antibody for RELT was incubated at 1:750 dilution in blocking buffer overnight at 4 °C. All other primary antibodies were diluted 1:1000 incubated at 1:1000 dilution in blocking buffer at 4 °C overnight and all blots were subsequently washed six times in TBS + 0.02% Tween-20 (TBS-T) for five minutes before a secondary antibody was incubated at 1:10,000 dilution for 45 minutes in blocking buffer. After washing with TBS-T three times for five minutes and TBS three times for five minutes, blots were imaged using either WestPico ECL (ThermoFisher) and Molecular Imager ChemiDoc XRS+ (BioRad, Hercules, CA) for HRP-conjugated secondary antibodies.

## Supporting information

Supplemental Information Cover PageF

## Supporting Information

Figure S-1. Quantification of LPS-induced secretion of cytokines IL6 and TNFα.

Figure S-2. Immunoblot of Hsc70 from immunoprecipitation control experiments indicating that Hsc70 is not being non-specifically eluted.

Table S-1. Complete listing of protein annotations in which at least one peptide was detected from any condition by in gel digestion of SDS-PAGE separated samples and in solution digested samples reported in this manuscript.

Table S-2. Complete listing of protein annotations in which at least one peptide was detected from any condition by in solution digest of isotype-controlled samples reported in this manuscript.

Table S-3. Full list of 292 high-confidence protein hits identified in both in gel and in solution digestions during this study.

Table S-4. Functional organization using DAVID of high-confidence proteins hits identified only in either LPS-treated or non-treated control conditions.

## Acknowledgements

JFR acknowledges the National Science Foundation (NSF-OIA-1738707) and NASA West Virginia EPSCoR Grant #NNX-15AK74A for support for this work. Some reagents were supplied by Dr. Ellis Bell (University of San Diego) through grant NSF-DUE-1726932. We thank Dr. Holly Cyphert (Marshall University) for providing access to her BioRad iMark Microplate Reader, Dr. Nadja Spitzer (Marshall University) for providing access to her BioRad GelDoc XRS+ and Dr. Robert Haltiwanger (University of Georgia) and his laboratory for providing invaluable technical assistance and manuscript advice, especially Mr. Steven Berardinelli. Lastly we thank Dr. Fikri Avci (University of Georgia, Athens, GA) for providing RAW264.7 cells.

## Conflict of Interest Disclosure

The authors declare no competing financial interest.

---

“for TOC only”

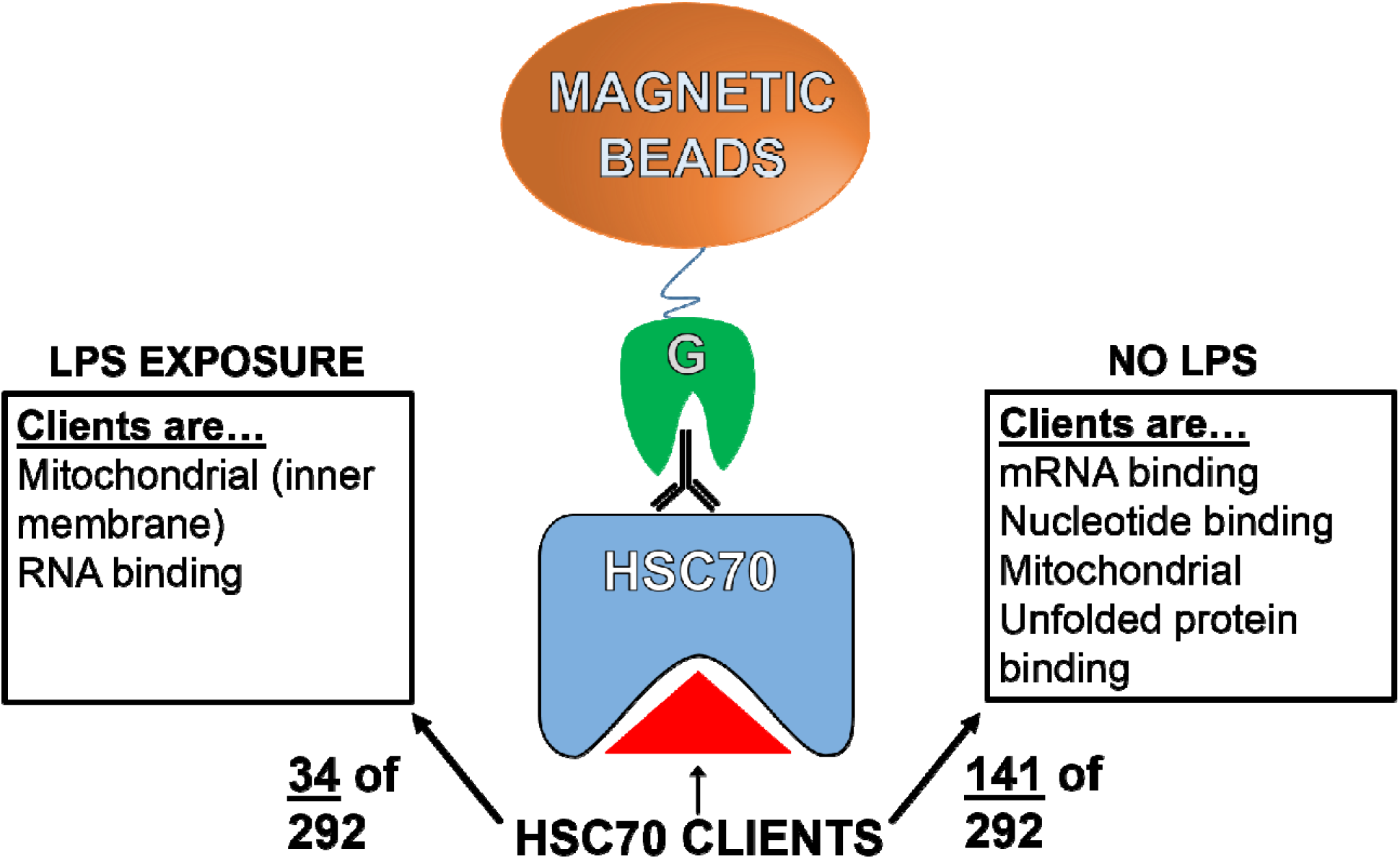

## Notes

### Competing Interest Statement

The authors have declared no competing interest.

