## Supplemental Information Cover PageF for "Lipopolysaccharide stimulation of RAW264.7 cells is a model for identifying novel clients of Hsc70"

**Table of Contents**

Figure S-1. Expression of the pro-inflammatory cytokines IL6 and TNFα in RAW264.7 cells grown in the absence or presence of 1 μg/mL LPS. Enzyme-linked immunosorbent assays (ELISAs) were purchased from R&D Systems and performed according to the manufacturer’s protocol.

Figure S-2. Hsc70 immunoblots of RAW264.7 cells from control conditions. Immunoprecipitation protocol was run identically to the in solution Hsc70 immunoprecipitation experiments. “Bead only” samples were non-conjugated Protein G-Dynabeads. “Isotype control” samples were Protein G-Dynabeads conjugated to a rat IgG control antibody.

Table S-1. Annotation of all proteins in which at least one peptide was detected in Byonic searches of in gel-digested Hsc70 samples. Each experimental condition is indicated and number of unique peptides for each protein are listed in the corresponding column. The percent coverage of the protein sequence is indicated in parentheses.

Table S-2. Annotation of all proteins in which at least one peptide was detected in Byonic searches of in solution-digested Hsc70 samples. Each experimental condition is indicated and number of unique peptides for each protein are listed in the corresponding column. The percent coverage of the protein sequence is indicated in parentheses.

Table S-3. Listing of protein annotations for high-confidence protein hits which were detected in both an in gel and in solution digestion. The annotations are organized into clusters based function as determined by DAVID. The DAVID enrichment score for each cluster is indicated as well as the gene ontology (GO) or Kyoto Encyclopedia of Genes and Genomes (KEGG) pathway entry for each cluster.

Table S-4. Listing of protein annotations for high-confidence protein hits grouped by proteins identified in either LPS-treated cells or non-treated control cells. The annotations are organized into clusters based function as determined by DAVID. The DAVID enrichment score for each cluster is indicated as well as the gene ontology (GO) or Kyoto Encyclopedia of Genes and Genomes (KEGG) pathway entry for each cluster.
