## Supplemental Figure 1 for "Lipopolysaccharide stimulation of RAW264.7 cells is a model for identifying novel clients of Hsc70"

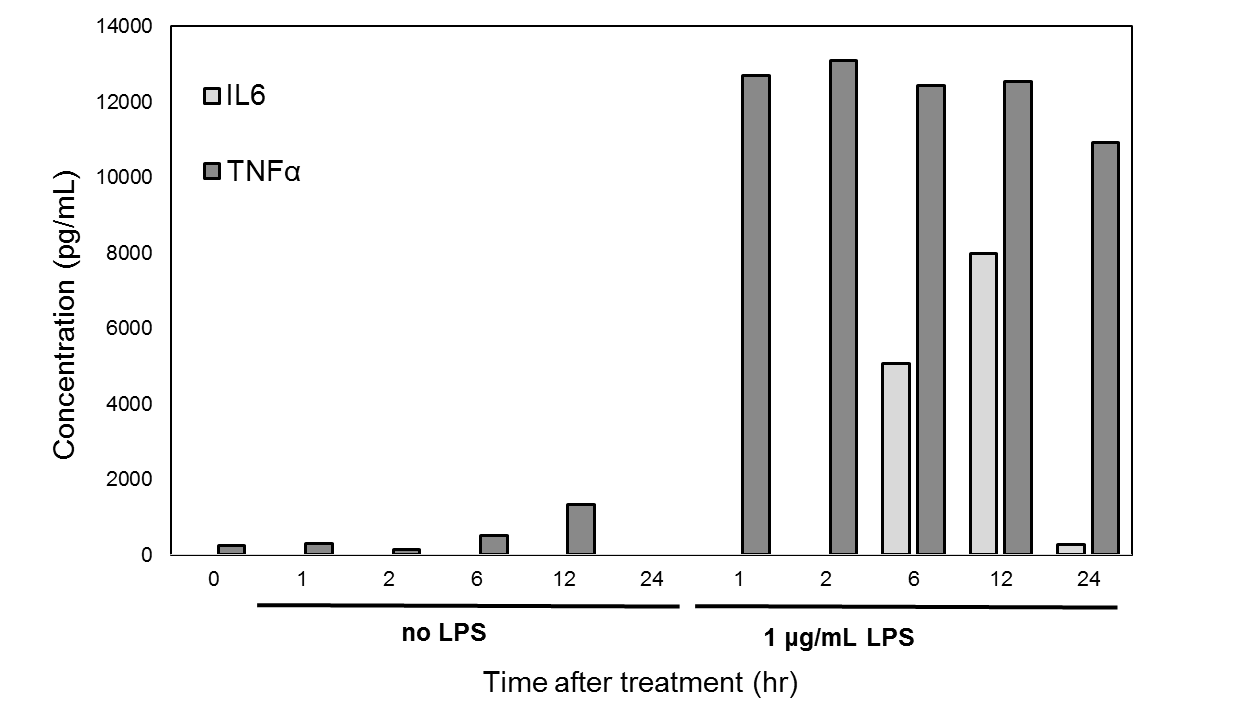

Figure S-1. Expression of the pro-inflammatory cytokines IL6 and TNFα in RAW264.7 cells grown in the absence or presence of 1 μg/mL LPS. Enzyme-linked immunosorbent assays (ELISAs) were purchased from R&D Systems and performed according to the manufacturer’s protocol.
