## Supplemental Figure 2 for "Lipopolysaccharide stimulation of RAW264.7 cells is a model for identifying novel clients of Hsc70"

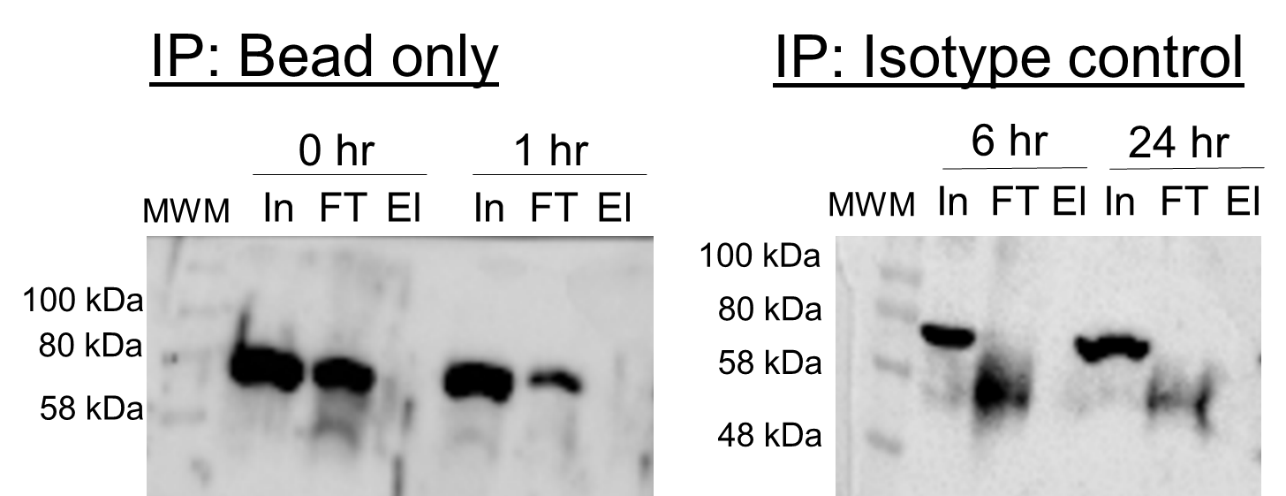

Figure S-2. Hsc70 immunoblots of RAW264.7 cells from control conditions. Immunoprecipitation protocol was run identically to the in solution Hsc70 immunoprecipitation experiments. “Bead only” samples were non-conjugated Protein G-Dynabeads. “Isotype control” samples were Protein G-Dynabeads conjugated to a rat IgG control antibody.
